# Empowering Student Authorship in Synthetic Biology

**DOI:** 10.1101/2024.03.30.587442

**Authors:** Louis A. Roberts, Natalie G. Farny

**Affiliations:** Department of Biology and Biotechnology, Worcester Polytechnic Institute, Worcester, MA USA; Program in Bioinformatics and Computational Biology, Worcester Polytechnic Institute, Worcester MA USA

**Author notes:** **Correspondence:** Louis A. Roberts, Natalie G. Farny.

**Keywords:** synthetic biology, higher education, peer review, authorship, primary literature

## Abstract

Women and racial minorities are underrepresented in the synthetic biology community. Developing a scholarly identity by engaging in a scientific community through writing and communication is an important component for STEM retention, particularly for underrepresented individuals. Several excellent pedagogical tools have been developed to teach scientific literacy and to measure competency in reading and interpreting scientific literature. However, fewer tools exist to measure learning gains with respect to writing, or that teach the more abstract processes of peer review and scientific publishing, which are essential for developing scholarly identity and publication currency. Here we describe our approach to teaching scientific writing and publishing to undergraduate students within a synthetic biology course. Using gold standard practices in project-based learning, we created a writing project in which students became experts in a specific application area of synthetic biology with relevance to an important global problem or challenge. To measure learning gains associated with our learning outcomes, we adapted and expanded the Student Attitudes, Abilities, and Beliefs (SAAB) concept inventory to include additional questions about the process of scientific writing, authorship, and peer review. Our results suggest the project-based approach was effective in achieving the learning objectives with respect to writing and peer reviewed publication, and resulted in high student satisfaction and student self-reported learning gains. We propose that these educational practices will contribute directly to the development of scientific identity of undergraduate students as synthetic biologists, and will be useful in creating a more diverse synthetic biology research enterprise.

## 1 Introduction

As a discipline that spans biology and engineering, synthetic biology tends to reflect a gender distribution among students and faculty that is closer to engineering disciplines than life sciences disciplines. In 2022, women made up 56.6% of life science doctorates awarded, but only 27% of engineering doctorates awarded (National Center for Science and Engineering Statistics (NCSES), 2023). Few formal studies have been conducted specific to synthetic biology; however, in biological and biomedical engineering departments across the U.S., women make up approximately 24.7% of faculty(American Society for Engineering Education, 2020). Student data is similarly scarce, but the limited information available suggests a similar skew in interest in synthetic biology by students. For a 2016 offering of “Principles of Synthetic Biology” as a massively open online course (MOOC) offered on edX, over 80% of the 11,768 people that engaged in the course identified as male (Anderson et al., 2019). Data on the representation of racial minorities in synthetic biology is even more paltry and is likely to reflect national populations of engineering students and faculty, suggesting low participation and retention of underrepresented students and faculty (American Society for Engineering Education, 2020; National Center for Science and Engineering Statistics (NCSES), 2023).

Developing a scholarly scientific identity is an important component for STEM engagement and retention, particularly for underrepresented individuals. Scientific identity is defined as having a sense of belonging to a scientific community, and seeing oneself as a valid member of that community (Carlone & Johnson, 2007). Many studies have suggested that developing a scientific identity through various educational, scientific, and professional development activities leads to greater STEM retention across the board, and can be particularly important in retaining diverse participants (Burt et al., 2023; Hernandez et al., 2017; Perez et al., 2014). Students’ self-assessed ability to communicate like a scientist – to speak with and to write to others within one’s discipline using a collectively accepted vocabulary and style – was shown to be a key indicator of future retention as a researcher within a given discipline (Cameron et al., 2020). These results suggest explicitly engaging undergraduate students in developing a scholarly identity through scientific communication skills may result in increased STEM retention. Thus, undergraduate coursework that promotes scientific communication skills will be an essential part of developing a more diverse pipeline of synthetic biologists.

Increasing students’ skills in scientific communication is among the core competencies identified in the latest recommendations for undergraduate life sciences education (American Association for the Advancement of Science, 2011). Several excellent tools have been developed to teach scientific literacy and to measure competency in reading and interpreting primary scientific literature (Goudsouzian & Hsu, 2023; Hoskins, 2008; Hoskins et al., 2007, 2011; Krufka et al., 2020). Scientific writing, which creates a tangible physical product to support a student’s scientific identity, has been proposed as an effective method of generative learning (also known as science writing heuristic and write-to-learn approaches) (Cronje et al., 2013; Reynolds et al., 2012). There are practices that engage students in authentic authorship experiences with publishable products (Burks & Chumchal, 2009; Giuliano et al., 2019; Guilford, 2001), though none have been described in the field of synthetic biology. Fewer tools exist to measure learning gains with respect to writing, or that teach the more abstract processes of peer review and scientific publishing (McDowell et al., 2019), and more development is needed in these areas(McDowell et al., 2022). Demystifying authorship and publication processes was shown to improve student learning outcomes and foster a greater sense of scientific identity among graduate students (Sletto et al., 2020). Similarly, increases in scientific identity were observed in undergraduate students that generated and analyzed their own data in course-based research experiences (Cooper et al., 2020; Roberts & Shell, 2023). The application of writing based approaches that incorporate authentic authorship experiences such as preprinting and peer review are thus of great importance in the development of scientific identities in undergraduate students.

Here we describe our approach to teaching scientific writing and publishing to undergraduate students within a synthetic biology course. Using gold standard practices in project-based learning (Larmer et al., 2015), we created a writing project in which students became experts in a specific application area of synthetic biology with relevance to an important global problem or challenge. To measure learning gains associated with our learning outcomes, we adapted and expanded the Student Attitudes, Abilities and Beliefs (SAAB) (Hoskins et al., 2011) assessment tool to include additional questions about the process of scientific writing, authorship, and peer review (which we refer to as SAAB-W). Students generated brief review articles that were publicly posted in the university digital archive. Our results suggest the project-based approach was effective in achieving the learning objectives with respect to writing and peer reviewed publication, and resulted in high student satisfaction and student self-reported learning gains. We propose that these educational practices will contribute directly to the development of scientific identity of undergraduate students as synthetic biologists, and will be useful in creating a more diverse synthetic biology research enterprise.

## 2 Materials and Methods

### 2.1 Course information

BB4260: Synthetic Biology is a seven-week upper-level elective course that meets for four contact hours per week (28 total contact hours). The intended student audience is upper level (junior or senior) undergraduate students, and early (first or second year) graduate students. The course is centered around analysis of primary literature in multiple areas of synthetic biology (e.g. genetic circuits, health applications, environmental applications, biocontainment, directed evolution), from the birth of the discipline in the year 2000 (Elowitz & Leibler, 2000; Gardner et al., 2000) through today. Thirty-one students enrolled in the course. Most students were juniors or seniors, and were life sciences majors and/or minors (Biology and Biotechnology, Biochemistry, Biomedical Engineering, and/or Bioinformatics and Computational Biology). Students self-identified their genders as 24 females and 7 males. There is no assigned textbook for the course; all assigned reading materials are open access journal articles. Credit is awarded in the course for participation in class discussions and group work (25%), in-class collaborative quizzes (40%), a final exam (10%) and the group writing project that is the focus of this research (25%). Learning outcomes (LOs) for the course as a whole and the project specifically (Table 1) were presented to students in the course and project syllabi, respectively.

**Table 1.**
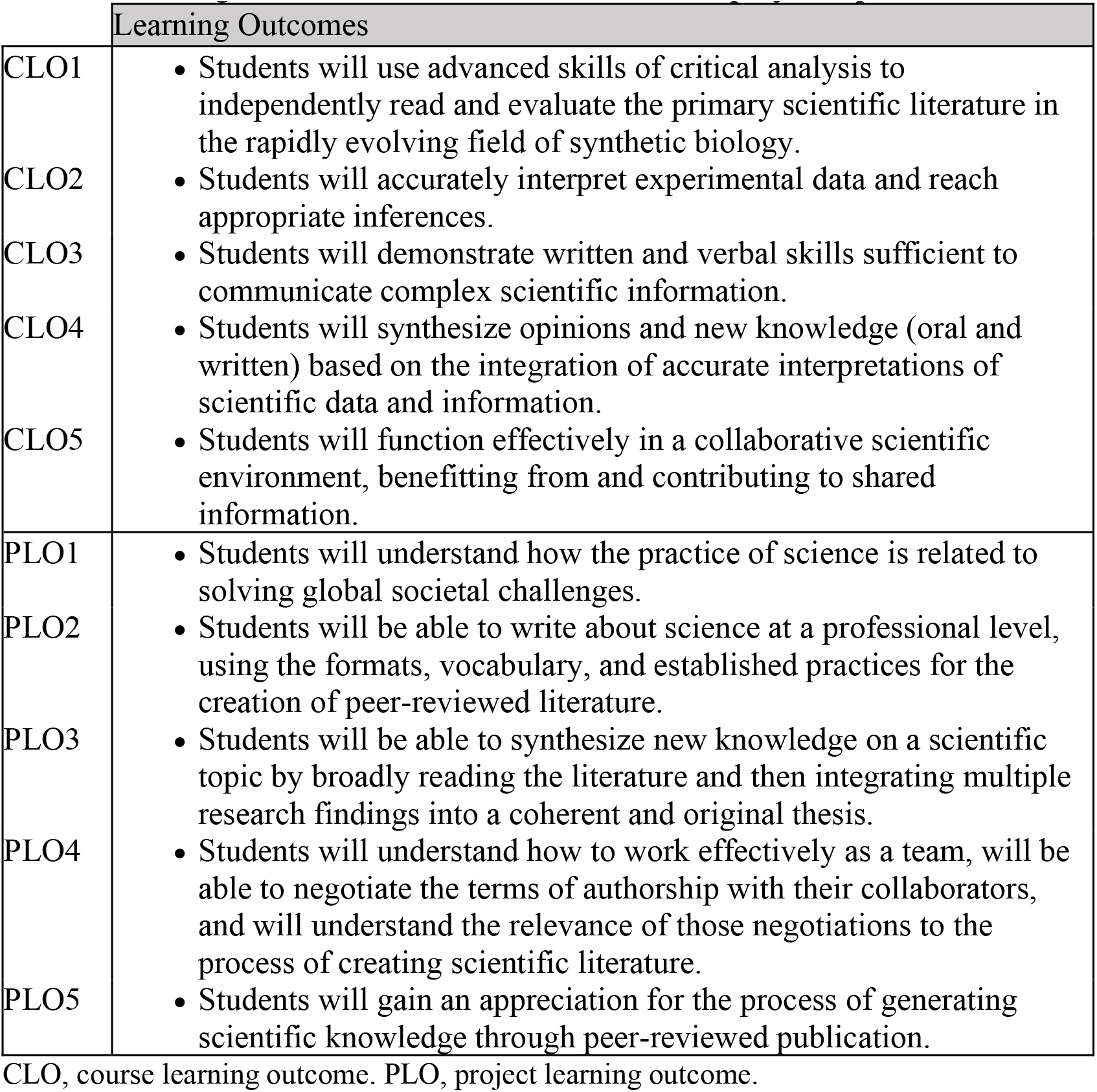
Learning outcomes for the entire course and the project experience.

### 2.2 Student Concept Inventory Survey

Students received multiple surveys throughout the course. The study protocol was reviewed and approved by the WPI Institutional Review Board (IRB-23-0611). The SAAB-W concept inventory (Table 2) pre-test was delivered to all students in the first week of the course using Qualtrics software. To maintain student anonymity and yet be able to match specific student pre- and post-test learning gains (LGs), students were asked to select a 4-digit code that was not revealed to the faculty. Students entered this code when taking both the pre- and post-test surveys. Surveys in which matching codes were available were included in the analysis (n=24). SAAB-W concept inventory data is anonymous and was not used for student assessment or grading. The survey administered to the students had 38 questions across the eight categories. When initially reviewing the survey and data, we noticed that one question on authorship (“The person who spends the most time writing (e.g. writes more of the article) receives most of the credit for a publication.”; reverse-scored) yielded an unusually large negative learning gain (−0.69). We decided to redact this result when presenting the data across the eight categories, and in relation to CLOs and PLOs. We feel this question and the results obtained are important when considered in the context of authorship, and use it as a valuable data point in that context as presented in the results (i.e., Figure 5).

**Table 2.** Concept inventory (SAAB-W) questions used for pre-/post-project student surveys.

| Question # | Question Prompt | Mapped LOs |
| --- | --- | --- |
| **1. Decoding Primary Literature** |  |  |
| 1-1 | The scientific literature is difficult to understand. (R) | CLO1,2 |
| 1-2 | When I see scientific journal articles, it looks like an unfamiliar language to me. (R) | CLO1,2 |
| 1-3 | I am not intimidated by the scientific language in journal articles. | CLO1 |
| 1-4 | I am confident in my ability to critically review scientific literature. | PLO3, CLO1,2 |
| 1-5 | I am comfortable defending my ideas about experiments. | PLO3, CLO2 |
| **2. Interpreting Data** |  |  |
| 2-1 | If I see data in a table, it is easy for me to understand what it means. | CLO2 |
| 2-2 | If I am shown data (graphs, tables, charts), I am confident that I can figure out what it means. | CLO2 |
| 2-3 | It is easy for me to relate the results of a single experiment to the big picture. | PLO1,3; CLO1,2 |
| 2-4 | Understanding the scales, axes, and legends for a graph or chart are essential for drawing any conclusions from experimental data. | PLO3; CLO1,3 |
| **3. Active Reading** |  |  |
| 3-1 | I could make a simple diagram that provides an overview of an entire experiment. | PLO3; CLO2,3 |
| 3-2 | If I am assigned to read a scientific paper, I typically read the methods section to understand how the data were collected. | CLO1 |
| 3-3 | I do not know how to design a good experiment. (R) | CLO4 |
| 3-4 | The way that you display your data can affect whether or not people believe it. | CLO4 |
| 3-5 | When I read a scientific article, I typically start at the beginning (abstract) and read straight through each section in order to the end (references). (R) | CLO1,2 |
| **4. Visualization** |  |  |
| 4-1 | When I read scientific information, I usually look carefully at the associated figures and tables. | CLO2 |
| 4-2 | When I read scientific material it is easy for me to visualize the experiments that were done. | CLO2 |
| 4-3 | If I look at data presented in a paper, I can visualize the method that produced the data. | CLO2 |
| 4-4 | When I read a paper, I have a clear sense of what physically went on in a lab to produce the results and information I am reading. | CLO2 |
| **5. Thinking Like a Scientist** |  |  |
| 5-1 | After I read a scientific paper, I don't think I could explain it to somebody else. (R) | PLO2,3; CLO3 |
| 5-2 | I am confident I could read a scientific paper and explain it to another person. | PLO2,3; CLO3 |
| 5-3 | I enjoy thinking of additional experiments when I read scientific papers. | PLO3; CLO4 |
| 5-4 | I accept the information about science presented in newspaper articles without challenging it (R). | PLO3 |
| 5-5 | I understand why experiments have controls. | CLO2 |
| **6. Scientific writing** |  |  |
| 6-1 | I can accurately summarize the content of a scientific paper in writing. | PLO2; CLO3 |
| 6-2 | I can combine observations from several papers to reach a broader conclusion. | PLO2,3; CLO1,4 |
| 6-3 | If asked to write a scientific article, I would be uncertain about how to begin that process. (R) | PLO2; CLO3,4 |
| 6-4 | It is difficult for me to draw connections between different research articles. (R) | PLO2,3; CLO1,4 |
| 6-5 | I can readily identify and summarize the key result of an experiment from an article. | PLO2; CLO1,2,3 |
| 6-6 | I know how and when to cite articles within a piece of scientific writing. | PLO2; CLO3 |
| 6-7 | I know how to locate scientific papers that are relevant to a specific topic. | PLO2 |
| **7. The process of scientific publishing** |  |  |
| 7-1 | I have a good understanding of how scientific papers are produced. | PLO4,5; CLO3 |
| 7-2 | I know how credit is assigned to authors within a list of authors on a publication. | PLO4; CLO5 |
| 7-3 | The person who spends the most time writing (e.g. writes more of the article) receives most of the credit for a publication. (R) | PLO4,5; CLO5 |
| 7-4 | I understand the process of peer review. | PLO5 |
| 7-5 | I understand the purpose of peer review. | PLO5 |
| **8. Global Relevance of SynBio Research** |  |  |
| 8-1 | I can identify why the authors of a paper believe their research is important. | PLO1; CLO1 |
| 8-2 | It is unclear to me why anyone would want to do synthetic biology research. (R) | PLO1; CLO1 |
| 8-3 | I am able to connect research objectives from a specific set of experiments to a broader societal goal. | PLO1; CLO1,4 |

Survey questions were grouped by topic and mapped to CLOs and PLOs. 28/31 students (90%) participated in these surveys. All surveys for which matching coded “pre” and “post” copies were available (n=24) were scored using the following five-point scale: strongly agree = 5, agree = 4, neither agree nor disagree = 3, disagree = 2 and strongly disagree = 1. Scores were inverted for questions marked (R).

### 2.3 Student Feedback Surveys

Students were asked at both the midpoint (Supplementary Figure S4) and the end (Supplementary Figure S6) of the project to provide feedback about their own contributions to the project, as well as their perceptions of contributions by their teammates. These surveys were a required element of the course and therefore were not anonymous to the course faculty. Students used a point system to express their opinions of the relative contributions they and teammates made to the project. Each student was also asked to use the CRediT contributor roles categories (Holcombe, 2019) to indicate the specific contributions they made to the project, as well as the relative effort they contributed (as a leader, collaborator, supporting, or no role). These surveys did contribute to student assessment and grading.

The university’s required student course report survey (Supplementary Table S2) was administered to all students using Class Climate software, according to university protocols. Students were given dedicated time during the class period to fill out the survey, during which the instructor left the room. These surveys are anonymous, and results of these surveys are not revealed to instructors until after final grades are submitted.

## 3 Results

### 3.1 Creating the Writing Project and Defining Project Learning Outcomes

When our synthetic biology course was created in 2021, we followed the principles of backwards course design to develop the course learning outcomes (Table 1). We utilized the same five course learning outcomes for the second offering in 2023. Inspired and supported by funding and expertise from our university’s EMPOwER (Engaging More Powerfully Openly with Educational Resources) program (Open Educational Resources (OERs) @ WPI, 2022), we set out to engage undergraduates with the primary literature to a greater depth, with the goal of stimulating their identity as scientists. We created a discrete writing project (worth 25% of the course grade) through which the students would engage with primary literature as purveyors and contributors. We next defined five project learning outcomes the writing project would fulfill (Table 1). We anchored the project to the 17 Sustainable Development Goals (SDGs) (UN DESA, 2023) as defined by the United Nations to reflect the global nature and societal implications of impactful science (PLO1). We believed it was vital for the students to become very familiar with established conventions and vocabulary utilized by the primary literature, so their writing would reflect the same tenor and tone (PLO2). Within the confines of a seven-week term and course format without a laboratory, we envisioned their contribution to the literature would be via authoring forward-facing review articles summarizing current knowledge, making connections across disciplines, and proposing new solutions synthetic biology may offer (PLO3). We emphasized the collaborative nature of writing such articles, which likely takes a different structure from the “divide and conquer” approach that can dominate other team-based class writing assignments. Thus, students were asked to engage in defining their authorship roles as guided by the CRediT taxonomy instrument, along with instruction in how authorship is defined and negotiated in the life sciences (PLO4). Students received explicit instruction on the path to publication, with a focus on the key role peer review plays in validating research contributions and knowledge generation. Students were then asked to exercise this information and empowered as reviewers of peer drafts in the course (PLO5).

### 3.2 Writing Project Workflow

We wanted to interweave the writing project with the course content, and provide the time required to satisfy the iterative nature of writing, reviewing, and feedback. We therefore utilized the entire seven-week term to carry out the writing project (Figure 2). The 31 students rank-ordered the SDGs based on their personal interest level in the Topic Survey (Supplementary Figure S1), and five project teams were formed based on their topic preferences (Week 1). Students were then directed to the primary literature to summarize the current state of knowledge relevant to their SGD and write a Topic Proposal (Week 2, Supplementary Figure S2) and complete an Annotated Bibliography assignment (Week 3, Supplementary Table S1). The instructor reviewed these to confirm the Topic Proposal clearly identified a roadblock to achieving an SGD that synthetic biology might address, and that the Annotated Bibliography was complete and aligned to their preliminary thesis statement. Student teams aligned their research efforts at this stage by creating a shared research summary (Supplementary Figure S3) which ensured students discussed and integrated the findings of their specific bibliographic annotations as a team. Next, student groups completed their initial rough drafts (Week 4), received feedback from the instructors, and refined these for a complete second draft (Week 5). The final versions of the article (including keywords, abstract, tables, and figures) were then submitted (Week 6). Peer review, in which students evaluated the work of other teams, was conducted using an instructor-provided template (Supplementary Figure S5) and a final round of instructor feedback was also conducted using the same peer review template (Week 7). Concept Inventory surveys (SAAB-W) were administered at the outset and upon completion of the writing project (as detailed below). Self- and peer evaluations were administered at the project mid-point and ending (Supplementary Figures S4, S6). Finally, the students completed a memorandum of understanding (MOU) regarding publication and authorship relevant to the deposit of their final product within DigitalWPI (Supplementary Figure S7).

**Figure 1.**
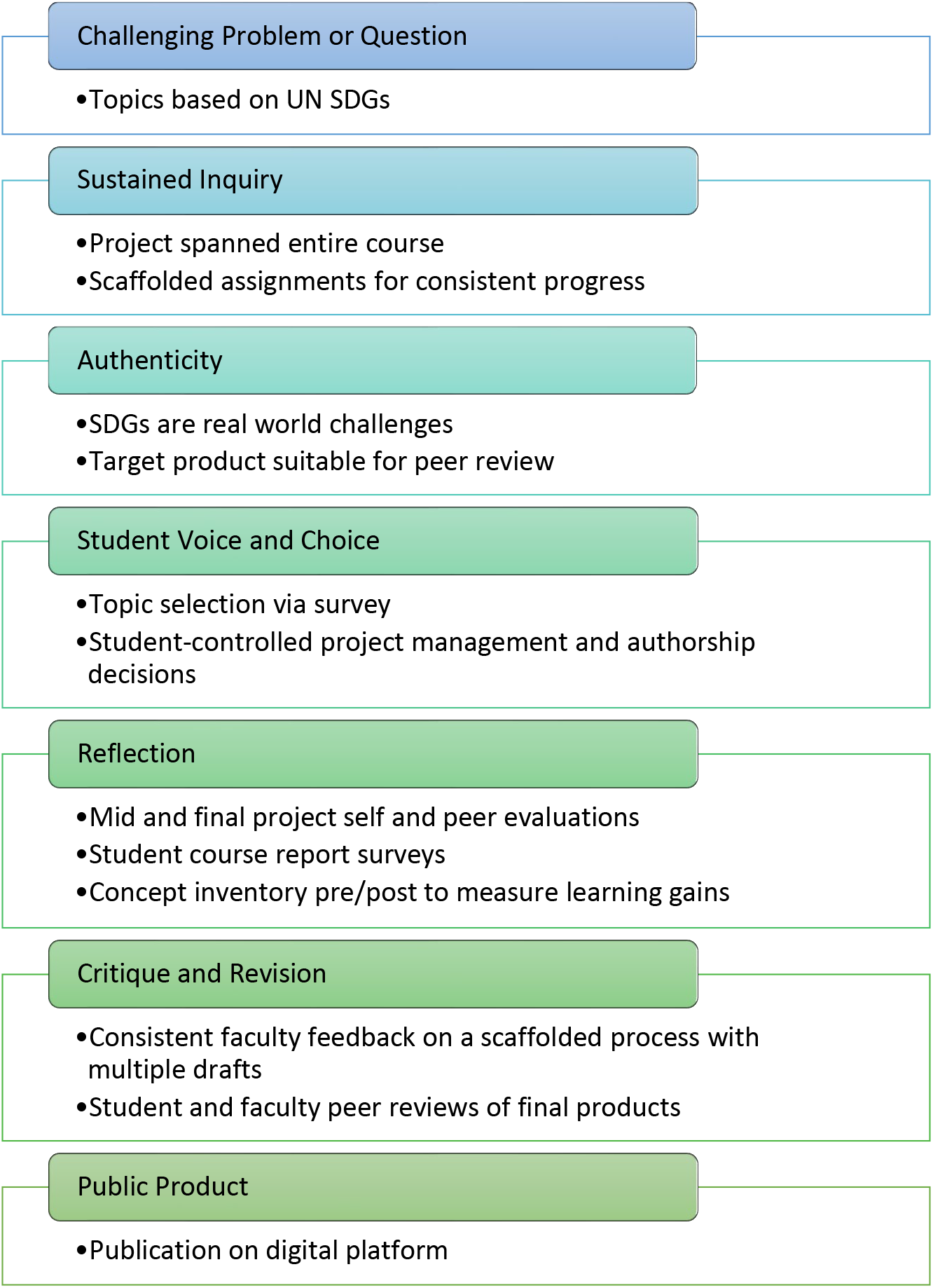
Project alignment with gold standard project-based learning design elements.

**Figure 2.**
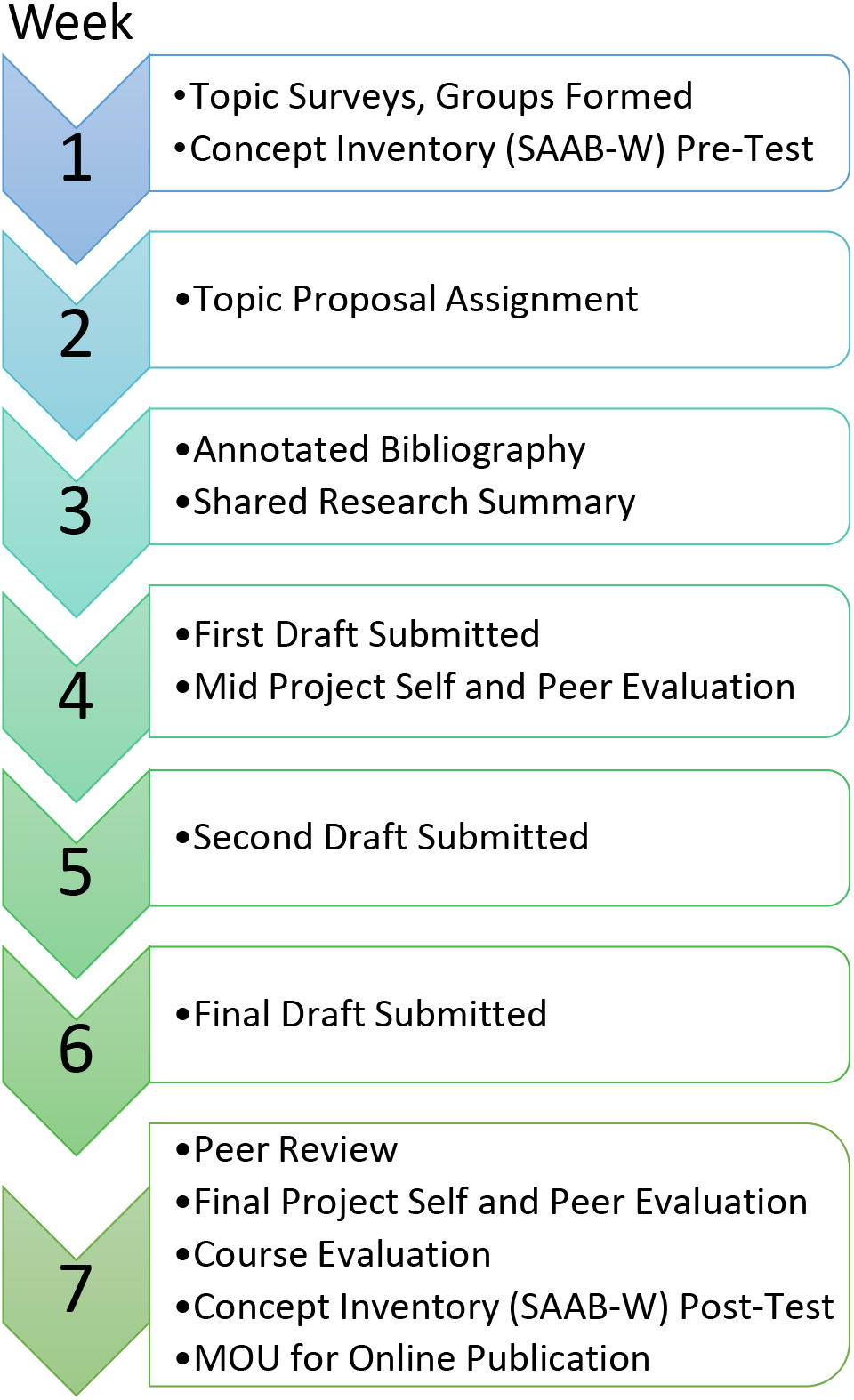
General timeline of assignments, activities, and assessments mapped onto the framework of our seven-week term.

### 3.3 Concept Inventory Development and Results

We adapted the Student Attitudes, Abilities, and Beliefs (SAAB) concept inventory (Hoskins et al., 2011; Krufka et al., 2020) as a starting point for assessing student learning gains (LGs) through their participation in the writing project. We used the Survey of factors 1-5 (Decoding the Primary Literature, Interpreting Data, Active Reading, Visualization, and Thinking Like a Scientist) to assess LGs in reading and critically analyzing the existing primary literature. Given that our PLOs reflect engaging students in generating new literature, we extended the concept inventory to include Scientific Writing, Process of Scientific Publishing, and Global Relevance of Synthetic Biology Research factors. We developed these questions by consulting with our university’s Writing Center director; the specific questions for each factor are shown in Table 2. Our concept inventory, which we have called SAAB-W, was administered just prior to introducing students to the writing project in Week 1, and then again after completing the writing project in Week 7.

The overall average LG reported for each of the eight factor categories is shown in Figure 3A. Positive LGs were observed for all categories, with the largest LGs for Process of Scientific Publishing (category 7; 0.94), Visualization (category 4; 0.85), and Scientific Writing (category 6; 0.75). Active Reading (category 3) showed the smallest LG (0.12). We then mapped each question within the eight categories to CLOs and PLOs, and plotted the average LG for each LO (Fig. 3B-C). Positive LGs > 0.4 were seen for all CLOs and PLOs. CLO5 (Students will function effectively in a collaborative scientific environment, benefitting from and contributing to shared information) and PLOs 4 and 5 (4: Students will understand how to work effectively as a team, will be able to negotiate the terms of authorship with their collaborators, and will understand the relevance of those negotiations to the process of creating scientific literature. 5: Students will gain an appreciation for the process of generating scientific knowledge through peer-reviewed publication.) showed the greatest LGs of 1.34, 1.23, and 1.12, respectively. Taken together, these results show students reported positive learning gains in each of the eight categories, five course learning outcomes, and five project learning outcomes. Given the direct link between the writing project and its specific set of learning outcomes (PLOs), we interpret these results to mean the writing project positively affected student learning as intended, and at a minimum did not impede learning in course outcomes nor in the standard SAAB factors designed to assess student learning in reading primary literature.

**Figure 3.**
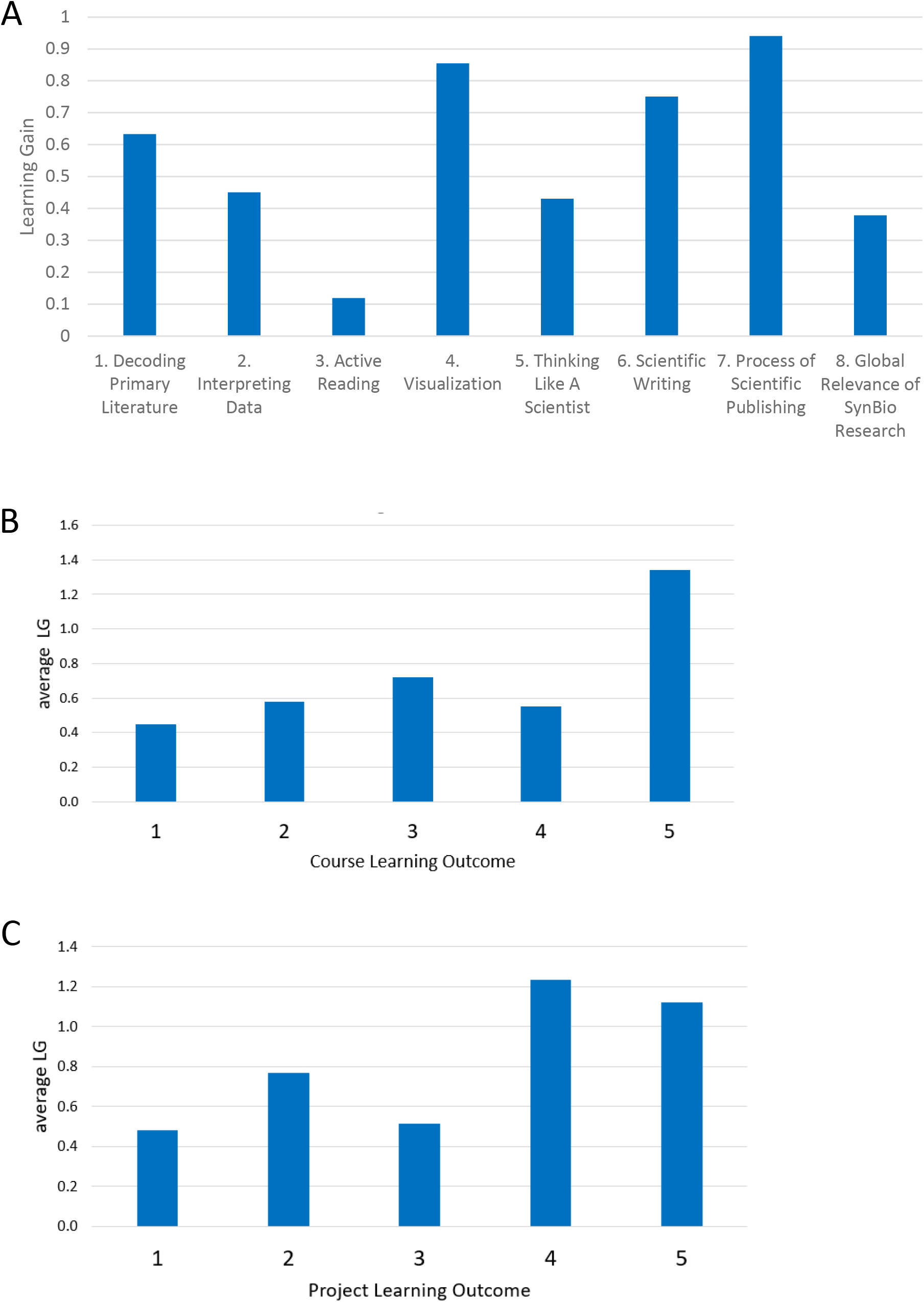
Learning gains in relation to SAAB-W concept inventory categories and learning outcomes. A) LGs for questions within each of the eight categories were averaged. B) LG averages vs. CLOs. C) LG averages vs. PLOs. For these analyses, question 7-3 was eliminated as described in the methods section. Analysis including question 7-3 is shown in Supplementary Figure S8.

The concept inventory results for each question are visualized in Figure 4. For each category, the raw pre and post scores are shown for the 24 student-matched surveys (Fig. 4A), along with the LG for each question (Fig. 4B). For the original SAAB instrument, which focuses on reading and analyzing the primary literature, consistent and notable positive LGs > 0.5 were obtained for all questions within Decoding the Primary Literature (category 1; five questions) and Visualization (category 4; four questions). Within these categories students reported entering the project near the midpoint of agreement/disagreement with the statements (2 < score < 4), implying a moderate level of incoming familiarity with the concepts, which were positively affected by completing the writing project and participating in the course. A similar landscape is observed for Interpreting Data (category 2; four questions) with one exception where student self-assessment was already very high (4.57) and a slight negative LG was seen (-.07). Similarly, within Thinking Like a Scientist (category 5; five questions) the only negative LG obtained was for question 5-5 regarding experimental controls (pre = 4.93, post = 4.85). Interestingly, only relatively small LGs (positive or negative) were reported for Active Reading (category 3; five questions); the two most positive LGs were related to understanding experimental design and workflow.

**Figure 4.**
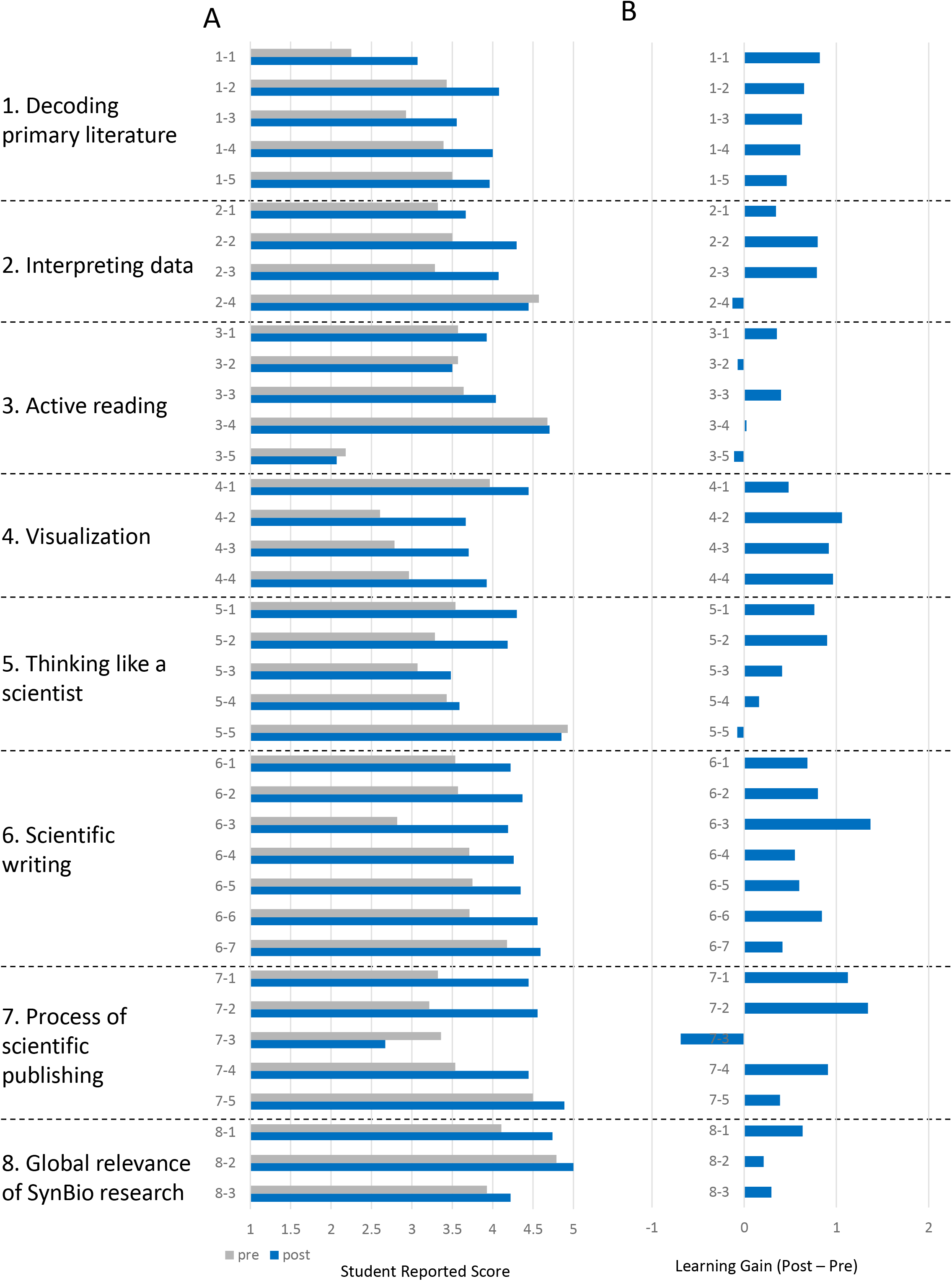
Students self-reported learning gains (n=24). A) Student pre- and post-project responses to the concept inventory survey questions from Table 2. B) Learning gains= (post – pre).

Focusing on responses to the questions we developed to specifically report on the writing project on student learning (Figure 4; categories 6-8), we achieved consistent and notably very positive LGs >0.4 for all questions in Scientific Writing (category 6; seven questions), and strong LGs within the Process of Scientific Publishing (category 7; four questions). The highest LG =1.37 across the 38 questions we posed was obtained for question 6-3, which explicitly focuses on how to initiate writing a scientific article. Given that writing a scientific article is a key PLO as well as a tangible product of the writing project, this data point validates our design and approach to promote student learning in the process of making new knowledge contributions to the literature. Positive LGs were further observed for all three questions in Global Relevance of SynBio Research (category 8). The students’ self-assessed opinion on entering the writing project was already in strong agreement that synthetic biology research is globally valuable (pretest value >3.9) and was likely a factor in why they registered for the course. Thus, the modest increases in LGs are understandable. In sum, the extended concept inventory survey and its data effectively gauged and reported student learning and validated our approach to engage students in exploring and creating primary literature.

### 3.4 Exploring Authorship Roles and Definitions

Emerging from our analysis of the data from the concept inventory was the surprising result of sharply negative LG (−0.69) for Question 7-3. The prompt states “The person who spends the most time writing (e.g., writes more of the article) receives most of the credit for a publication”. We reverse-scored this item because within the life sciences, credit is not typically assigned by volume of writing (for example, a corresponding author may write most of the manuscript in some cases, or in other cases the first author(s) may do so). We used the contributor roles taxonomy (CRediT) model (Holcombe, 2019) to guide our discussions with the class and explain the concept that authorship roles are determined by a range of contributions from intellectual contributions to technical execution to financial support to supervision. Inspecting the results at a higher resolution (Figure 5A), we noticed that 12 of 24 students from matched surveys had greater agreement post-project that writing volume is the determining factor; ten students did not change their belief; and four students has greater disagreement post-project that writing volume is the main factor when defining authorship roles. The negative LG suggests many students may have become more entrenched in their belief the amount of writing weighs heavily in determining authorship credit, contrary to our expectations. Undergraduates completing team writing assignments in other courses and projects experience a linkage between writing contribution and grades; equal grading for the product of a group assignment implicitly or explicitly demands equal contributions in writing. Course-based group writing therefore exists in a different context than creating peer-reviewed literature, and our students have significantly more experience with the former. Given the nature of the writing project in which a written document is the main product, alternative avenues of contributing (i.e., performing experiments, providing financial support, etc.) were either abstract or not relevant. We then considered whether other data we acquired might shed light on how students defined what credit for authorship means, and the relative values they placed on forms of credit (e.g., grades vs. authorship).

**Figure 5.**
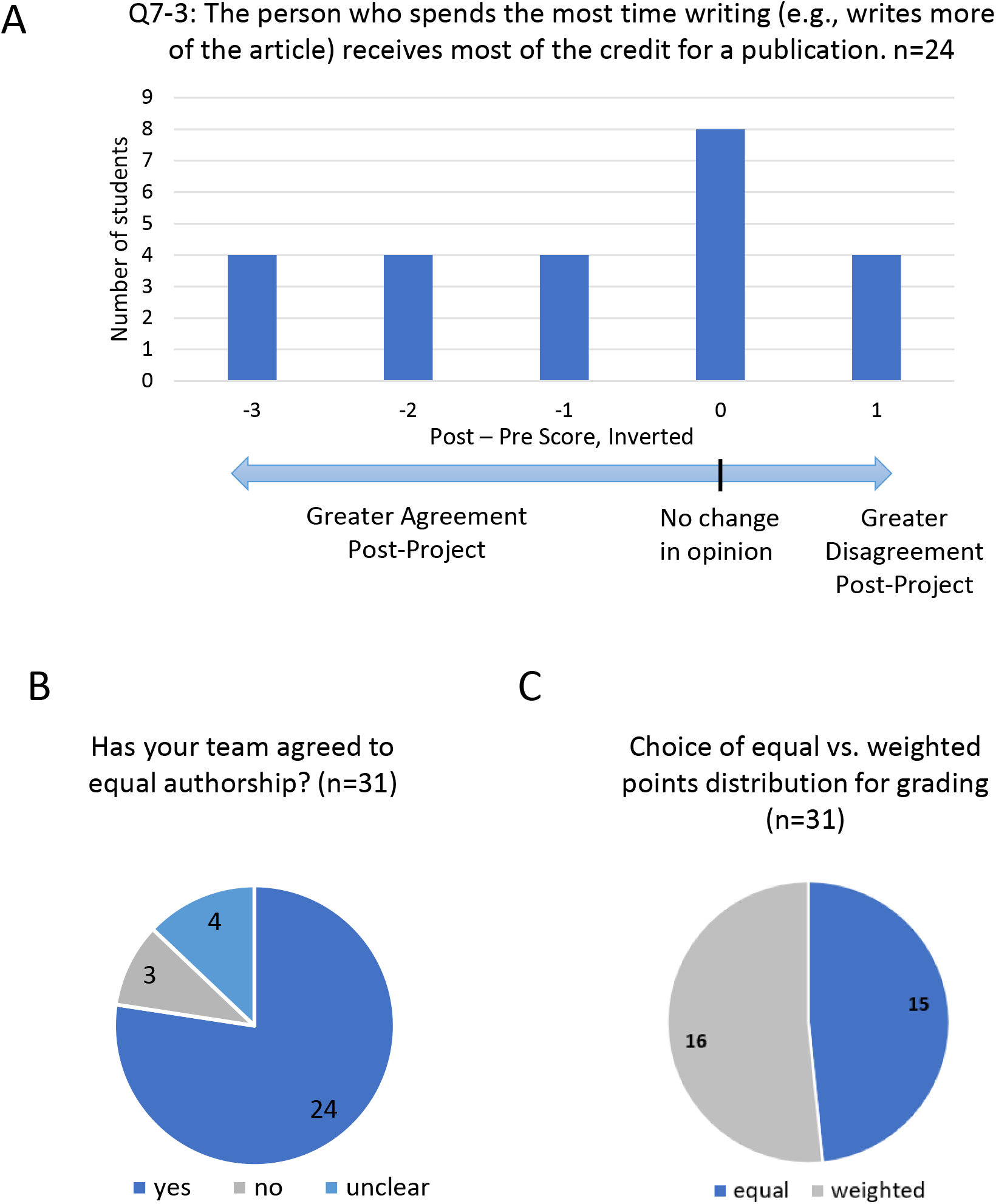
Exploring authorship and grading credit. A) Binned analysis of changes in response to Q7-3 (post-project score minus pre-project score, inverted). B) Negotiating authorship for non-peer reviewed submission to library. C) Weighting of points for writing project grade.

A final project evaluation survey was administered (Supplementary Fig. S6, required and thus not anonymous; n=31) which asked students whether their group had negotiated equal versus weighted authorship with respect to their submission to the university library open access repository DigitalWPI. Students were separately asked how they believed points should be awarded for grading the assignment; it was made clear in the syllabus and MOU (Supplementary Fig. S7) that these two issues were distinct, and that consent to authorship in DigitalWPI was unrelated to the final grade for the course. These data are aligned by groups in Table 3. 24 students responded their groups had agreed to equal authorship (Figure 5B), including all members of Groups 2 and 4. Groups 1 and 3 had three respondents who were unclear on the authorship discussion but commented they did feel equal authorship was warranted. The three students who responded “no” to the question on whether equal authorship was agreed upon were in Group 5. This group negotiated authorship order collegially, based on administration and work/writing invested in the project as stated in their comments. One student explicitly referred to applying CRediT taxonomy to determine author order. Thus, the responses strongly suggest all five groups negotiated authorship order for submission into DigitalWPI, and for this purpose four groups (1-4) agreed to equal authorship roles.

**Table 3.** Final Project Survey- Defining Authorship and Contributions.

| Group | # students | Negotiated DigitalWPI authorship? | Equal DigitalWPI authorship? | Same group and individual grade? | Equal weighting of contribution points? |
| --- | --- | --- | --- | --- | --- |
| 1 | 6 | Yes | Yes | 6 yes | 3 yes<br>3 no |
| 2 | 6 | Yes | Yes | 6 yes | 6 yes<br>0 no |
| 3 | 7 | Yes | Yes | 7 yes | 1 yes<br>6 no |
| 4 | 6 | Yes | Yes | 4 yes; 1 higher group; 1 higher individual | 4 yes<br>2 no |
| 5 | 6 | Yes | No | 5 yes; 1 higher group | 0 yes<br>6 no |

Student responses on final team project evaluation survey (n=31).

The final project survey also asked students to confidentially distribute 30 “contribution” points across their group; approximately half (16/31) chose an unequal distribution of points (Figure 5C and Table 3), including all six students in Group 5 who agreed to weighted authorship. Group 2 had all members uniformly distribute points; the remaining ten students who chose unequal weightings for contributions were scattered across the three remaining groups. We asked the students to state the grades they believe the team project and they deserve. 28/31 reported the same grade for the team and themselves; two recommended a higher grade for the team, and one responded with a higher individual grade. Taken together, most students were content with equally shared authorship and equal project grades, even though several of these students recommended an unequal allotment of points for contributions to the project.

The final project survey also asked students to identify their contribution level (i.e., lead, collaborative, supporting) in each of the 14 CRediT categories, and whether they intended to continue revising the article to submit for peer-reviewed publication. Student responses to the five CRediT roles most relevant to the course writing project (Conceptualization, Investigation, Project Administration, Writing-Original Draft; Writing-Review and Editing) are shown in Figure 6A. Students could define their role as lead, collaborator, supporting, or no role. For all five CRediT categories, a notable majority of students chose their role as collaborator. For four of the categories, ≤5 students selected lead or supporting as their role. Notably, 10 respondents (30%) saw their role as lead for the Writing-Review and Editing CRediT category, with at least one falling in each student group. Given this role is largely unfolding late in the project period (in comparison to Writing-Original Draft), it may reflect the reality of the contextual, temporal nature of undergraduate coursework as much as explicitly defined student roles in the project.

**Figure 6.**
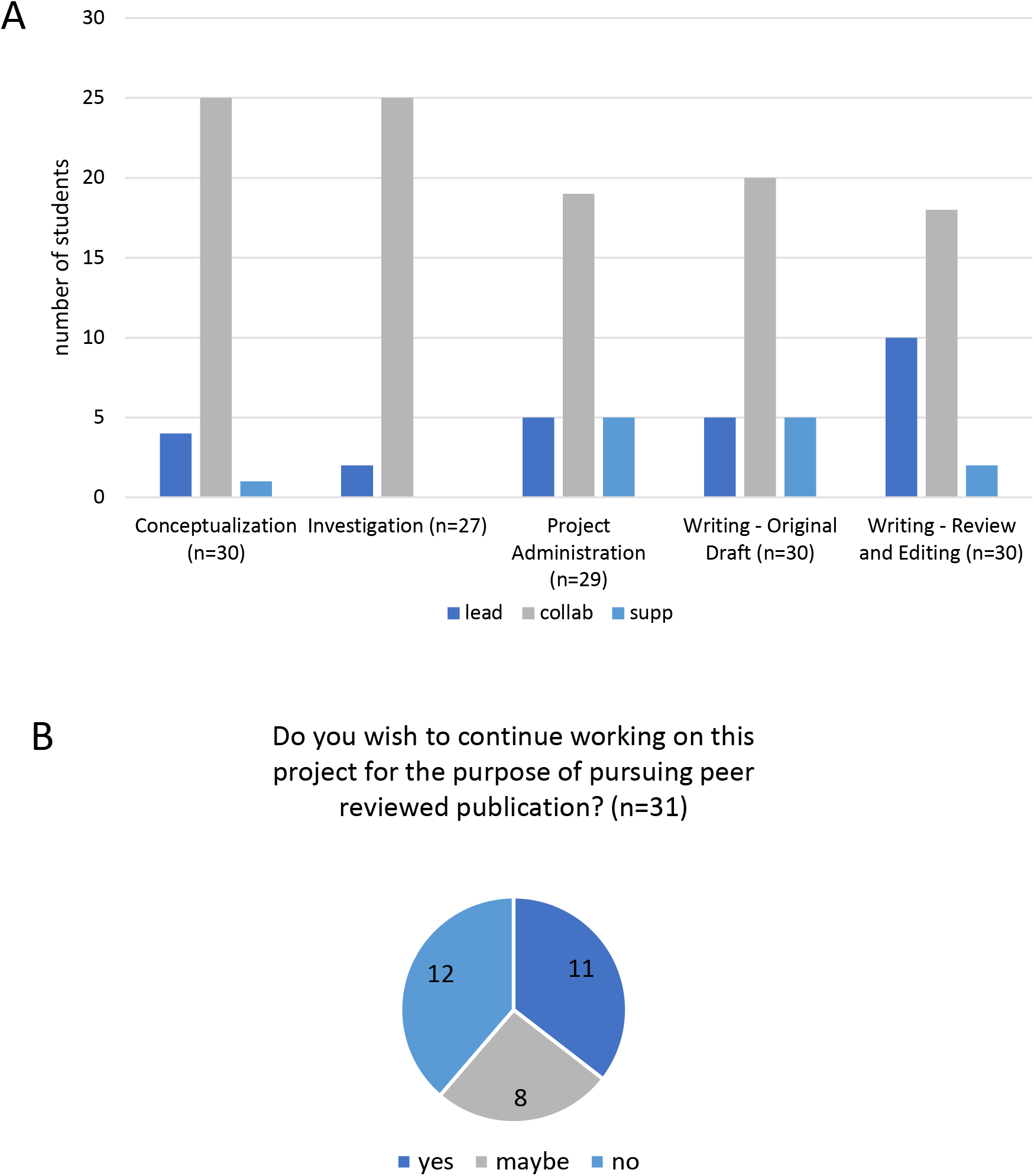
Students’ perceptions of their contributions to the project. A) Students reported contributions to the project, based on CRediT contributorship categories. Lead, leading role in this task. Collab, collaborative role in this task. Supp, supporting role in this task. Not all CRediT categories applied to the project or the student authors, thus only categories in which >80% of students selected a response are reported. B) Students’ responses to the question about their interest in continuing to work on the project.

The minimum requirement for the final project document was a version suitable for inclusion in the DigitalWPI open access collection we created entitled “Synthetic Biology for Global Good” (Farny & Roberts, 2023). The writing project was designed to produce original work consistent with the standards of a peer-reviewed publication. We instructed the students to write in the style of a “Forum” piece for *Trends in Biotechnology*, and posted several examples of such articles on the course website. All five groups generated articles for DigitalWPI, thus meeting the requirement for the course. We extended an optional opportunity for students to continue to refine their articles for submission to a peer-reviewed journal. The final project evaluation survey asked the students their relative interest level engaging in the publication process beyond the course period (Figure 6B). Of the 31 students, 11 expressed strong interest in continuing; 12 stated they did not wish to participate; eight responded “maybe”. As several students were graduating seniors and the currency was authorship rather than a course grade, we were pleased with the number of students willing to pursue publication.

### 3.5 Course level survey results

Our synthetic biology course has been offered four times to date, twice with the current CLOs, and two prior offerings in 2019 and 2020 as a “Capstone” course, offered with small class sizes (∼15 students) and rotating topics. The writing project was first deployed in 2023, replacing the former group assignment to present a research paper to the class. There were no other significant changes to the course learning outcomes, other assignments, workflow, or course website, and the instructor (Dr. Farny) was the same for all offerings. This class has historically received strongly positive student course evaluation scores as measured by WPI’s course evaluation survey, completed by students at the end of the term. To assess the impact of the project experience on student reported course satisfaction, student course evaluation scores were compared across all four course offerings (Table 4). The 2023 offering, which included the new writing project, retained the very high scores previous versions of the course received. We conclude the writing project did not detract from students’ perceptions of the course’s quality and value.

**Table 4.** Student Course Evaluations.

| Question Text | 2019<br>(n=12) | 2020<br>(n=16) | 2021<br>(n=11) | 2023<br>(n=23) |
| --- | --- | --- | --- | --- |
| 1. My overall rating of the quality of this course is | 5.0 | 4.9 | 4.8 | 4.9 |
| 3. The educational value of the assigned work was | 4.8 | 4.8 | 4.8 | 4.8 |
| 4. The instructor's organization of the course was | 4.9 | 4.8 | 4.9 | 4.8 |
| 7. Relative to other college courses I have taken, the amount I learned from the course was | 4.7 | 4.6 | 4.7 | 4.6 |
| 8. Relative to other college courses I have taken, the intellectual challenge presented by the course was | 4.4 | 3.9 | 4.6 | 4.4 |
| 10. Relative to other college courses I have taken, the instructor stimulated my interest in the subject matter | 4.7 | 4.6 | 4.6 | 4.8 |

Selected questions from the university’s standardized required course evaluation. Questions 1, 3 and 4 are on a scale of 1 (very poor) to 5 (excellent). Questions 7, 8, and 10 are on a scale of 1 (much less) to 5 (much more). The number of student responses is given for each question.

The final question on the writing project survey gave the students the option to provide general comments on their team and the writing project, and 12 students chose to provide comments. To visualize the responses, we created a Word Cloud (Figure 7) to see which words students used to describe the team-based writing project. While abstract and non-quantitative in nature, we value the time and thoughtfulness students invest in providing feedback. The visualization is helpful to gain an understanding of how the students perceived the writing project as a component of the synthetic biology course.

**Figure 7.**
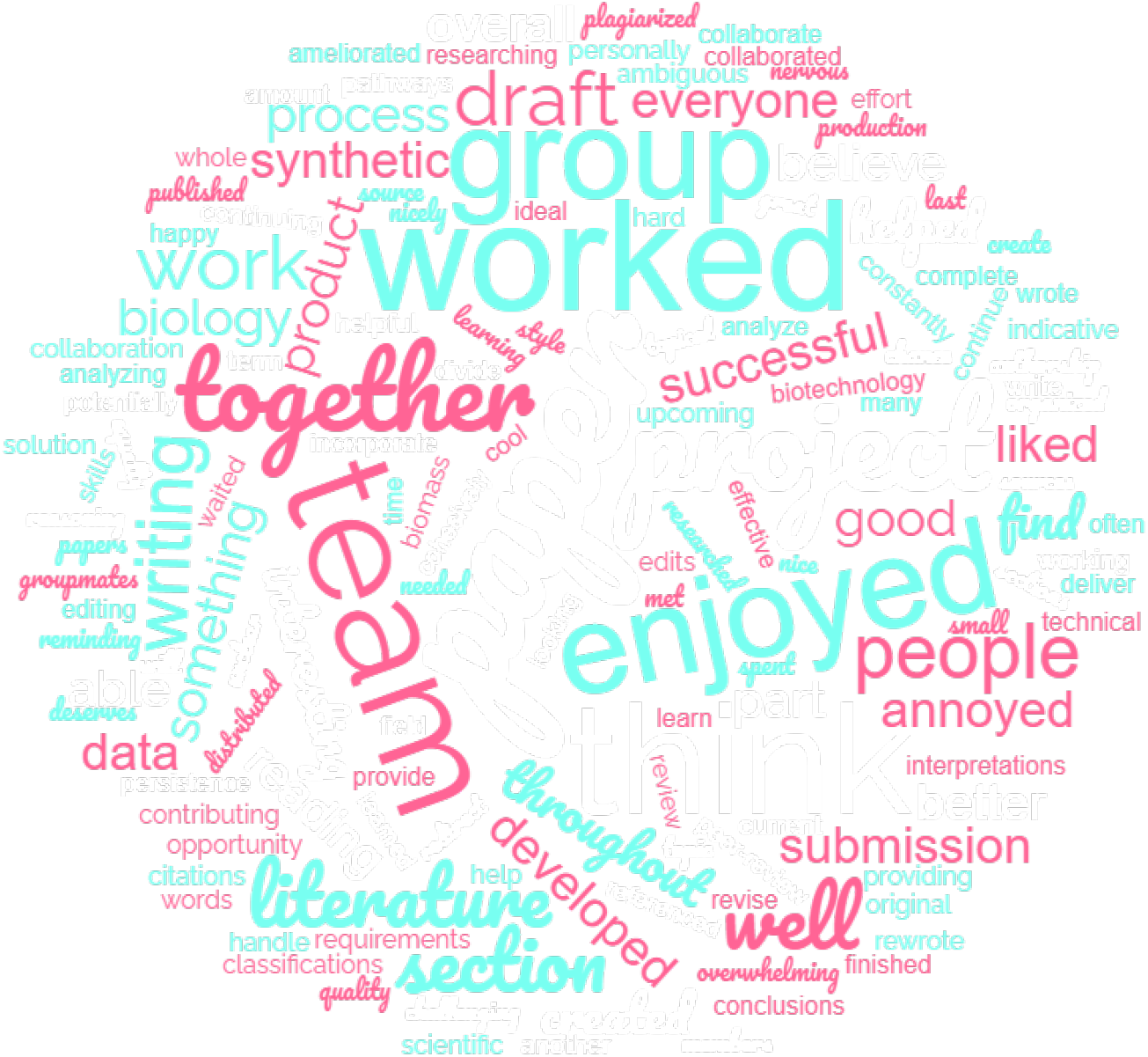
Word Cloud of student comments from writing project survey (2023).

Both the university course evaluations and the Final Project Self and Peer Evaluation surveys provided a place for students to comment on the course and assignments. Representative comments focused on the writing project included:

“This course has been a huge help in improving my practical skills related to reading and understanding scientific literature, all while providing very interesting topics to learn from in doing so.” - Student Author 1

“I really enjoyed this project as whole. I liked researching a current issue and looking into how synthetic biology could provide a solution. It was also very interesting to learn about the peer-review process as this will likely be very helpful while working in the biotechnology field.” -Student Author 2

“I think that our project really developed nicely from our original draft. I think our topic was a little challenging but we created something interesting and very topical. I enjoyed the opportunity to write a review style paper, which is often something I did not get the chance to do. I also really enjoyed learning about the authorship classifications. Overall, I think I learned a lot from the project and it was nice to collaborate with a group of people who I hadn’t worked with before!” -Student Author 3

“When the section of the second draft I wrote had ambiguous writing and referenced data with unclear reasoning and interpretations, I rewrote my section and further researched the literature to find more ideal papers with better data and conclusions. Throughout the writing process, I really developed my skills in reading and analyzing scientific literature as well as technical writing.” - Student Author 4

## 4 Discussion

We were motivated to develop new teaching approaches that would increase the diversity of the synthetic biology community by providing students the opportunity to engage with and contribute to the primary literature, thereby supporting the development of their scholarly identities. Funded by a university grant program (EMPOwER), we created a new writing project with five specific learning outcomes, and embedded the project in an existing Synthetic Biology course. A unique feature of the project is students were instructed in how to write in the style of a primary literature article (mini-review) that was ultimately published on our university library’s open access platform (DigitalWPI), and participate in the peer review process. To ascertain if the learning outcomes for the project and the class were achieved from the student perspective, we used standard assessments (the SAAB-W inventory and university course evaluations). We worked with our university’s Writing Center to expand the original SAAB inventory (which centers around reading and evaluating the primary literature) to include self-assessment questions for three new categories-Scientific Writing, Process of Scientific Publishing, and Global Relevance of SynBio Research. We created a collection entitled “Synthetic Biology for Global Good, Volume I” (Farny & Roberts, 2023) featuring five articles written by 31 students. We will use this resource as a model for future student project work, as well as assigned reading to introduce complex topics within the course. We plan to expand upon the collection in future iterations of the course.

Our data support that our approach to teach undergraduates scientific writing and publishing by immersing them in an authentic authorship experience was successful. All five writing project PLOs show sizable learning gains as measured by our expanded SAAB-W survey. All 38 questions were mappable to project and/or course learning outcomes. Thus, we were satisfied that the additional categories and questions we created for this survey were adequate to ascertain how the students assessed their experience of the writing process within the project in the context of the course. We are considering optimizations to the questions we added, and the SAAB-W survey as a whole, perhaps reducing the overall number of questions to streamline data collection and minimizing survey burnout among the students participating in the course.

Our students (mostly juniors and seniors) entered the course possessing confidence in reading journal articles. Moving beyond reading the primary literature, significant learning gains in knowledge of how articles are written, how authorship is defined in our field, and both the process and the purpose of peer review were observed. This knowledge has demystified for these students a critical component of success in STEM fields, that of contributing to scientific knowledge through publication. In line with our overarching goal, the broader data in the literature suggests that such efforts lead to increased development of scholarly identity and, therefore, better STEM engagement and retention, particularly for underrepresented and minoritized students (Burt et al., 2023; Hernandez et al., 2017; McDowell et al., 2022; Perez et al., 2014).

One unexpected observation that emerged when analyzing the survey data is how students perceive authorship relative to the specific context of writing. We expected students would leave the course acknowledging authorship roles for the primary literature are defined by intellectual input rather than writing output. A negative LG was actually obtained on the SAAB-W survey (Question 7-3). We reflected upon this surprising result, and observed that most student groups either assumed or ignored equal intellectual input, and instead assessed contributions based on effort, time, and volume of writing. Our university is immersed in project-based learning (Larmer et al., 2015), which frequently uses student teams/groups to complete project objectives that often include a final written report. Our students experienced this writing project in the context of a course, and seemingly applied their existing experience and framework to define contributions. Interestingly, students and groups most often chose to share authorship equally, using grading (via weighted points distribution) as a way to acknowledge contribution level. Our data may indicate that this additional level of development, that of assessing relative intellectual contributions and shedding pre-conceived notions of what constitutes a significant contribution, is still in process for students at this stage and was not completely addressed by our pedagogical approach.

The writing project was specifically designed for and embedded in our upper-level Synthetic Biology course, and we were curious what effects the project may have on the overall student learning experience in the class. We mapped the student self-reported LGs from the SAAB-W survey back onto the course learning outcomes, and examined our university-wide formal course evaluations over several offerings. While the SAAB-W survey is specifically tailored to the writing project and thus not explicitly informed by the CLOs, the compatibility of the project and the Synthetic Biology course informed our decision to embed the project within this class. Students reported positive LGs for all five CLOs, which supports the view that carrying out the writing project did not detract from, and possibly enhanced, student learning within the course as a whole. The university course evaluations maintain the very high scores that had been achieved in previous offerings, and we were encouraged by a slight elevation in the score for stimulating interest in the subject matter (4.8 up from 4.6 in the previous version of the class). These surveys indicate students perceived learning and value from completing the course with the embedded writing project we designed.

A significant challenge in a seven-week term is the time available to undertake iterative writing with multiple drafting-feedback-refine cycles, particularly for feedback and peer review. For instance, while all students engaged in the peer review process and comments from peer review were collated and returned to groups at the end of the course, teams did not have time to respond to peer reviews by incorporating edits or writing a rebuttal. With respect to the writing project timeline as implemented, the five groups were able to meet on schedule all milestones for the project. At the level of resolution of individual students within the groups, over half chose an unequal weighting of points, often supported by comments regarding contribution level. This may or may not relate to the ability of individual students to meet self-imposed or instructor-set deadlines, and we will consider adding a specific question to our Final Project Self and Peer Evaluation Survey to gain insight in future offerings of the project. On the flip side, the course instructor(s) also have a short window of time to guide the groups to define their topics, provide feedback on two drafts, and organize and then disseminate the in-class peer review. While constrained by the condensed term schedule, we were fortunate the writing project utilized two faculty instructors to rapidly provide feedback to the five groups. The much more common course format (14-week semester) expands the timeframe for instructor feedback, and thus a second faculty member participating in the writing project may not be necessary. The Synthetic Biology class did have one graduate student teaching assistant (TA) who focused exclusively on the course and not on the writing project; we view in a standard course format a TA could contribute to reviewing the manuscript drafts and organizing the student groups and peer review process.

Overall, student feedback about the project was largely positive, with most students expressing that the project was a valuable experience. Of the 31 students in the course, 24 self-identified as female and 7 self-identified as male. In the field of synthetic biology, which takes much inspiration from male-dominated engineering fields, women are in the minority as authors of the primary literature. This project opportunity allowed more women to develop scholarly identities as authors within this field, which is an important contribution to the diversification of synthetic biology. The feedback reflects the achievement of most of the key learning goals for the project, including the process of reading literature, technical writing, and the process of peer-reviewed authorship.

Consistent with our motivation to embrace open access publication of our student articles, all the teaching materials used for the writing project (including the SAAB-W inventory) are freely available in this article and the supplementary material. While our writing project was embedded in a Synthetic Biology lecture course, we envision the project timeline, methods, and tools will translate well into any upper-level life sciences STEM course. We hope our approach will inspire other educators to adopt and adapt these ideas to support their own students’ future STEM careers.

## Author Contributions

N.G.F. conceived of and designed the course and the project, and was the primary instructor for the course. L.A.R. performed data analysis and curation, and primarily wrote the first draft. Both authors contributed to design of the assessments, surveys, and course materials, provided feedback on student work, assembled figures, tables, and supplementary material, edited and finalized the manuscript, and obtained financial support.

## Acknowledgments

We are grateful to the entire WPI EMPOwER team for their critical support of this project, particularly Marja Bakermans and Lori Ostapowicz-Critz, and student feedback from Hannah Shell. We thank Ryan Madan, Director of the WPI Writing Center, for his guidance in helping to design the extended SAAB-W categories and questions.

## Supplementary Material

Supplementary Material including Supplementary Figures S1-S2 and Supplementary Tables S1-S8 accompany this manuscript and are available online.

## Data Availability Statement

The anonymized datasets for this study are available upon request to N.G.F., in accordance with IRB protocol limitations to protect student privacy.

## References

American Association for the Advancement of Science. (2011). Vision and change in undergraduate biology education: A view for the 21st century. https://www.aps.org/programs/education/undergrad/upload/Revised-Vision-and-Change-Final-Report.pdf

American Society for Engineering Education. (2020). Engineering and Engineering Technology by the Numbers 2019. https://ira.asee.org/wp-content/uploads/2021/02/Engineering-by-the-Numbers-FINAL-2021.pdf

Anderson, D. A., Jones, R. D., Arkin, A. P., & Weiss, R. (2019). Principles of synthetic biology: a MOOC for an emerging field. Synthetic Biology, 4(1), ysz010.

Burks, R. L., & Chumchal, M. M. (2009). To co-author or not to co-author: how to write, publish, and negotiate issues of authorship with undergraduate research students. Science Signaling, 2(94), tr3–tr3.

Burt, B. A., Stone, B. D., Motshubi, R., & Baber, L. D. (2023). STEM validation among underrepresented students: Leveraging insights from a STEM diversity program to broaden participation. Journal of Diversity in Higher Education, 16(1), 53.

Cameron, C., Lee, H. Y., Anderson, C. B., Trachtenberg, J., & Chang, S. (2020). The role of scientific communication in predicting science identity and research career intention. PloS One, 15(2), e0228197.

Carlone, H. B., & Johnson, A. (2007). Understanding the science experiences of successful women of color: Science identity as an analytic lens. Journal of Research in Science Teaching: The Official Journal of the National Association for Research in Science Teaching, 44(8), 1187–1218.

Cooper, K. M., Knope, M. L., Munstermann, M. J., & Brownell, S. E. (2020). Students who analyze their own data in a course-based undergraduate research experience (CURE) show gains in scientific identity and emotional ownership of research. Journal of Microbiology & Biology Education, 21(3), 10–1128.

Cronje, R., Murray, K., Rohlinger, S., & Wellnitz, T. (2013). Using the science writing heuristic to improve undergraduate writing in biology. International Journal of Science Education, 35(16), 2718–2731.

Elowitz, M. B., & Leibler, S. (2000). A synthetic oscillatory network of transcriptional regulators. Nature, 403(6767), 335–338.

Farny, N. G., & Roberts, L. A. (Eds.). (2023). Synthetic Biology for Global Good, Volume 1. Worcester Polytechnic Institute. https://digital.wpi.edu/concern/generic_works/8g84mq800?locale=en

Gardner, T. S., Cantor, C. R., & Collins, J. J. (2000). Construction of a genetic toggle switch in Escherichia coli. Nature, 403(6767), 339–342.

Giuliano, T., Skorinko, J. L. M., & Fallon, M. (2019). Engaging undergraduates in publishable research: Best practices. Frontiers in Psychology, 10, 482812.

Goudsouzian, L. K., & Hsu, J. L. (2023). Reading Primary Scientific Literature: Approaches for Teaching Students in the Undergraduate STEM Classroom. CBE—Life Sciences Education, 22(3), es3.

Guilford, W. H. (2001). Teaching peer review and the process of scientific writing. Advances in Physiology Education, 25(3), 167–175.

Hernandez, P. R., Bloodhart, B., Barnes, R. T., Adams, A. S., Clinton, S. M., Pollack, I., Godfrey, E., Burt, M., & Fischer, E. V. (2017). Promoting professional identity, motivation, and persistence: Benefits of an informal mentoring program for female undergraduate students. PloS One, 12(11), e0187531.

Holcombe, A. O. (2019). Contributorship, not authorship: Use CRediT to indicate who did what. Publications, 7(3), 48.

Hoskins, S. G. (2008). Using a paradigm shift to teach neurobiology and the nature of science—a CREATE-based approach. Journal of Undergraduate Neuroscience Education, 6(2), A40.

Hoskins, S. G., Lopatto, D., & Stevens, L. M. (2011). The CREATE approach to primary literature shifts undergraduates’ self-assessed ability to read and analyze journal articles, attitudes about science, and epistemological beliefs. CBE—Life Sciences Education, 10(4), 368–378.

Hoskins, S. G., Stevens, L. M., & Nehm, R. H. (2007). Selective use of the primary literature transforms the classroom into a virtual laboratory. Genetics, 176(3), 1381–1389.

Krufka, A., Kenyon, K., & Hoskins, S. (2020). A single, narrowly focused CREATE primary literature module evokes gains in genetics students’ self-efficacy and understanding of the research process. Journal of Microbiology & Biology Education, 21(1), 10–1128.

Larmer, J., Mergendoller, J., & Boss, S. (2015). Setting the standard for project based learning. Ascd.

McDowell, G. S., Fankhauser, S., Saderi, D., Balgopal, M., & Lijek, R. S. (2022). Use of preprint peer review to educate and enculturate science undergraduates. Learned Publishing, 3, 405–412.

McDowell, G. S., Knutsen, J. D., Graham, J. M., Oelker, S. K., & Lijek, R. S. (2019). Co-reviewing and ghostwriting by early-career researchers in the peer review of manuscripts. Elife, 8, e48425.

National Center for Science and Engineering Statistics (NCSES). (2023). Doctorate Recipients from U.S. Universities: 2022. https://ncses.nsf.gov/pubs/nsf24300/

Open Educational Resources (OERs) @ WPI. (2022). EMPOwER. https://wp.wpi.edu/open/

Perez, T., Cromley, J. G., & Kaplan, A. (2014). The role of identity development, values, and costs in college STEM retention. Journal of Educational Psychology, 106(1), 315.

Reynolds, J. A., Thaiss, C., Katkin, W., & Thompson Jr, R. J. (2012). Writing-to-learn in undergraduate science education: a community-based, conceptually driven approach. CBE—Life Sciences Education, 11(1), 17–25.

Roberts, L. A., & Shell, S. S. (2023). A research program-linked, course-based undergraduate research experience that allows undergraduates to participate in current research on mycobacterial gene regulation. Frontiers in Microbiology, 13, 1025250.

Sletto, B., Stiphany, K., Futrell Winslow, J., Roberts, A., Torrado, M., Reyes, A., Reyes, A., Yunda, J., Wirsching, C., & Choi, K. (2020). Demystifying academic writing in the doctoral program: Writing workshops, peer reviews, and scholarly identities. Planning Practice & Research, 35(3), 349–362.

UN DESA. (2023). The Sustainable Development Goals Report 2023: Special Edition - July 2023. https://desapublications.un.org/publications/sustainable-development-goals-report-2023-special-edition

