## Supplemental Figures and Tables for "Empowering Student Authorship in Synthetic Biology"

### 1 Supplementary Figures and Tables

### 1.1 Supplementary Figures

For our project "SynBio for Global Good", you will write about some area of research within synthetic biology that is contributing to one of the United Nations Sustainable Development Goals.

Please read about each of the goals by clicking on the goal, here: https://sdgs.un.org/goals.

Arguably, synthetic biology has something to offer toward each and every goal. What global challenge are you most interested in thinking about? Please rank the SDGs in your order of interest by dragging and dropping them to re-order them. Every effort will be made to put you into a group based on your first or second choice, but this may not be possible if there are too few students to form groups around a particular topic. We appreciate your flexibility. Thanks!

Please enter your name [text box]

Please rank order these SDGs according to your interests [drag-and-drop rank order]
1. No poverty
2. Zero hunger
3. Good health and well-being
4. Quality education
5. Gender equality
6. Clean water and sanitation
7. Affordable and clean energy
8. Decent work and economic growth
9. Industry, innovation, and infrastructure
10. Reduced inequalities
11. Sustainable cities and communities
12. Responsible consumption and production
13. Climate action
14. Life below water
15. Life on land
16. Peace, justice, and strong institutions
17. Partnerships for the goals

**Supplementary Figure S1.** Project Topic Survey. This survey was distributed using Qualtrics software.

**A. Team Introduction**
Please create a team roster with name, year, major, and email address for each student. (add rows to the table as needed)

| Name | Class year | Major | Email |
|---|---|---|---|

Give yourselves a fun team name!

**B. Authorship Considerations**
All students are expected to contribute to this project as part of the requirements of this course. However, some students may find they are more invested in bringing the project to publication than others (which will NOT affect your grade in either direction, aiming for *Trends in Biotechnology* is optional). In some groups, you may feel that folks contributed more or less equally – how then should authorship be attributed? Other groups may feel that some members contributed more significantly than others – how will you handle that? What about if some students want to stay engaged with this project past the end of the course, and others don't? How then might that change the authorship landscape?
Please have a frank conversation with your group about your thoughts and feelings on the topic of authorship. What you want to contribute and what you hope to achieve, personally and professionally, are all part of this conversation. In the space below, summarize your conversation, and come up with an action plan for managing authorship.
Please note that all students will do a "self-audit" using the CRediT taxomony of contributorship at both the midpoint and final team evaluations, and that the team action plan may be adjusted as we go forward.
________________________________________________________________________
____________________________________________

**C. Topic Ideas**
Which SDG has your group chosen? __________________________________
The SDGs are broad – you will need to narrow your topic significantly because you only have 1200 words! Brainstorm some specific ideas for topics relevant to this SDG here. You might want to use PubMed or Google Scholar to search for key words related your topic ideas and see what's out there. Please note that this is not a firm commitment at this stage, and sometimes topics evolve as you dig into the literature.
________________________________________________________________________
________________________________________________________________

Do you have any ideas for visual items (text boxes, figures, tables)? It may be hard to decide this before you zero in on a more focused topic, but jot down any initial thoughts here. Look through the example papers posted in the Project module for some ideas.
_______________________________________________________________
___________________________________________________________________

**D. Work Plan**
Who wants to do what? What needs to be done within the next few weeks in order to get to a solid first draft? Assign a role or roles for each team member for the next several weeks. I recommend that you choose a team captain or captains who will turn in deliverables and manage group documents, but you may opt to distribute these responsibilities if you prefer.
Outline your work plan below.
Please note that ALL team members MUST contribute 3 articles each to the annotated bibliography.
________________________________________________________________________
________________________________________________________________________
______________________________________________________________

**Supplementary Figure S2.** Topic Proposal Assignment Template. These assignment prompts were used by teams to define their specific project topics. One assignment was submitted per team.

**Shared Research Summary**

1) Talk with each other and share key points of your sources.

2) Write up a short summary of the main ideas you learned from each of the articles (from the annotated bibliography assignment) and think about themes that run across all the articles. In your summary, also integrate relevant information (terms, concepts) from the course readings/discussions.

3) List/describe areas where you think more research is necessary.

This can take the form of a paragraph, or it can take the form of a bulleted list. All of the information included in your summary should be details, data, support, etc., that are **relevant** to and **build** assignment 1. Use this opportunity to order/organize the information in a logical manner and integrate course terms and concepts--again to help you build toward assignment 1. This is also an opportunity to identify 1-2 figures, graphs, or text boxes you'd like to create that illustrates your important thesis or common theme.

Please INCLUDE the names of people who 1) shared their findings from their annotated bibliography, 2) contributed to the summary (discussion, writing, etc.), and 3) both.

One team member should email this summary to Prof. Farny before the end of class – copy all team members. If a team member is not present during class time to work with the team, they need to upload a second version where they integrate their information into the work that was completed during class (please highlight where text was added, thank you).

**Supplementary Figure S3.** Shared Research Summary Template.

The purpose of this evaluation is to understand how individuals are contributing to the group project at this stage, and to give you the opportunity to provide feedback on the function of the project team. Required and optional questions are noted.

[Required] Please enter your name.

[Required] In this first section, please use the space below to distribute a total of 30 points among your team members (including yourself) based on the quality and importance of each individual's contribution. Please be honest and objective as much as possible.

Use each field to put a team member's name, then the points you would like to assign to that team member. For example:

*Natalie Farny, 6 points.*

Please list yourself first.

Team member 1
Team member 2
Team member 3
Team member 4
Team member 5
Team member 6
Team member 7

[Optional] Please use this space to comment on any of the points assignments you made above.

[Required] Below are listed the CRediT Taxonomy categories. Please indicate your own contributions using the taxonomy, to this point in the project.
As a reminder, descriptions of these CRediT categories can be found at https://credit.niso.org/.

| | Lead contributor (more than equal contribution) | Collaborative contributor (about equal contributor) | Supporting contributor (less than equal contribution) | Did not contribute | not applicable |
|---|---|---|---|---|---|
| Conceptualization | ○ | ○ | ○ | ○ | ○ |
| Data curation | ○ | ○ | ○ | ○ | ○ |
| Formal analysis | ○ | ○ | ○ | ○ | ○ |
| Funding acquisition | ○ | ○ | ○ | ○ | ○ |
| Investigation | ○ | ○ | ○ | ○ | ○ |
| Methodology | ○ | ○ | ○ | ○ | ○ |
| Project administration | ○ | ○ | ○ | ○ | ○ |
| Resources | ○ | ○ | ○ | ○ | ○ |
| Software | ○ | ○ | ○ | ○ | ○ |
| Visualization | ○ | ○ | ○ | ○ | ○ |
| Validation | ○ | ○ | ○ | ○ | ○ |
| Supervision | ○ | ○ | ○ | ○ | ○ |
| Writing, original draft | ○ | ○ | ○ | ○ | ○ |
| Writing, review and editing | ○ | ○ | ○ | ○ | ○ |

[Optional] Please use this space to provide any other comments or thoughts about the team or the project more generally.

**Supplementary Figure S4.** Mid Project Self and Peer Evaluation Survey. Surveys were prepared and distributed using Qualtrics software.

**Peer Review**: [article title]

This review article by [author last names] describes... (one sentence topic description)

The authors' thesis/key observation/critical insight is... (try to re-state their thesis/main point/primary contribution)

**Summary of article content** (3 sentences – written objectively without criticism or judgement, just a statement of what you read)

**Strengths/positive feedback** (general, 2-3 sentences, this is where you can infuse your opinions)

**Weaknesses/areas for improvement** (general, 2-3 sentences, same here, it is appropriate in this section to offer opinions on what you read)

**Specific Comments** (can be bulleted lists)

**Major issues:** include specific suggestions for edits, areas where ideas are unclear, or sentences that don't read well or are ambiguous in their meaning. You would also include comments if you happen to disagree with the authors, or feel they are missing some key points or data from the literature (this type of comment is easier to do if you are an expert in the field and may not be possible at your career stage).

**Minor issues**: include typos, grammatical errors, formatting, etc in this last section. These are the least critical elements of the review and are often handled by the production editor, but it can be helpful to point them out if you notice them.

**Supplementary Figure S5.** Peer Review Template.

The purpose of this evaluation is to understand how individuals contributed to the group project, and to give you the opportunity to provide feedback on the function of the project team. Required and optional questions are noted.

Please consider the finished project, and your contributions as a whole, as you complete this survey.

[Required] Please enter your name.

[Required] In this first section, please use the space below to distribute a total of 30 points among your team members (including yourself) based on the quality and importance of each individual's contribution. Please be honest and objective as much as possible.

Use each field to put a team member's name, then the points you would like to assign to that team member. For example:

*Natalie Farny, 6 points.*

Please list yourself first.

Team member 1
Team member 2
Team member 3
Team member 4
Team member 5
Team member 6
Team member 7

[Optional] Please use this space to comment on any of the points assignments you made above.

[Required] Below are listed the CRediT Taxonomy categories. Please indicate your own contributions using the taxonomy, to this point in the project.
As a reminder, descriptions of these CRediT categories can be found at https://credit.niso.org/.

| | Lead contributor (more than equal contribution) | Collaborative contributor (about equal contributor) | Supporting contributor (less than equal contribution) | Did not contribute | not applicable |
|---|---|---|---|---|---|
| Conceptualization | ○ | ○ | ○ | ○ | ○ |
| Data curation | ○ | ○ | ○ | ○ | ○ |
| Formal analysis | ○ | ○ | ○ | ○ | ○ |
| Funding acquisition | ○ | ○ | ○ | ○ | ○ |
| Investigation | ○ | ○ | ○ | ○ | ○ |
| Methodology | ○ | ○ | ○ | ○ | ○ |
| Project administration | ○ | ○ | ○ | ○ | ○ |
| Resources | ○ | ○ | ○ | ○ | ○ |
| Software | ○ | ○ | ○ | ○ | ○ |
| Visualization | ○ | ○ | ○ | ○ | ○ |
| Validation | ○ | ○ | ○ | ○ | ○ |
| Supervision | ○ | ○ | ○ | ○ | ○ |
| Writing, original draft | ○ | ○ | ○ | ○ | ○ |
| Writing, review and editing | ○ | ○ | ○ | ○ | ○ |

**Supplementary Figure S6.** Final Project Self and Peer Evaluation Survey. Surveys were prepared and distributed using Qualtrics software.

**Supplementary Figure S6 Cont.**

[Required] Has your team agreed to equal authorship, at least for the purpose of this stage of submission to Digital WPI?

- ○ yes
- ○ no (please explain below) ______
- ○ authorship credit is unclear to me (please explain below) ______

[Required] Do you wish to continue working on this project for the purpose of pursuing peer reviewed publication?

- ○ Yes, definitely
- ○ Maybe, I need more information (explain below) ______
- ○ No, definitely not

[Required]. If you do not want to continue to work on the project, do you have any concerns about other group members continuing to do so? Please keep in mind, all team members will continue to be updated by email as to the status of any peer reviewed publication, even if they choose not to continue to contribute to the project, unless they specifically request to no longer be copied on such correspondence.

- ○ No questions or concerns
- ○ Yes, I have questions/concerns (explain below) ______

[Required] What grade do you feel ***your team*** deserves for this project?

- ○ A
- ○ B
- ○ C

[Required] What grade do you feel ***you personally*** deserve for this project?

- ○ A
- ○ B
- ○ C

[Optional] Please use this space to provide any other comments or thoughts about the team or the project more generally.

**MOU for Students and Faculty**
**Agreement to Contribute to Open Educational Resource**

I, ______________________________, agree to participate in the creation of ***Synthetic Biology for Global Good, Volume I***, an open educational resource, in collaboration with my professor, Natalie Farny. This work will comprise part of my coursework for BB4260: Synthetic Biology. This work will be made publicly available on the Digital WPI online repository.
I understand that inclusion of my work in the final collection is conditional upon my willingness to license my contributions under a CC-BY-NC license. This license allows re-users to distribute, remix, adapt, and build upon the material in any medium or format for noncommercial purposes only, and only so long as attribution is given to the creator. I have had the opportunity to review the description of this license at https://creativecommons.org/about/cclicenses/.
I understand that I have the right to request that my name and/or work be removed from the original text, or change the license on my contributions at any stage prior to publication.

Signed: ______________________________ Date: ___________________

I, Natalie Farny, agree to work with the student ______________________________ on the creation of ***Synthetic Biology for Global Good, Volume I***, an open educational resource, in partial completion of the course BB4260: Synthetic Biology, to be made publicly available on the Digital WPI online repository.
I commit to supporting ___________________ throughout this project, and ensuring they have the knowledge and resources they need to be an informed contributor.
I agree that the student may request that their name and/or work be removed from the original text or change the license on their contributions to this work at any stage prior to publication of the work.
I confirm that the student's decision to change the license they place on their work or to not participate in the public posting of this project will not impact on their course assessment. A student's decision to decline to participate in this agreement will result in the student's name being removed from the publicly available work.
Signed: _ ______________________ Date: __________________

**Supplementary Figure S7.** Memorandum of Understanding. Students and faculty signed an agreement about the terms of publication on the university's digital repository.

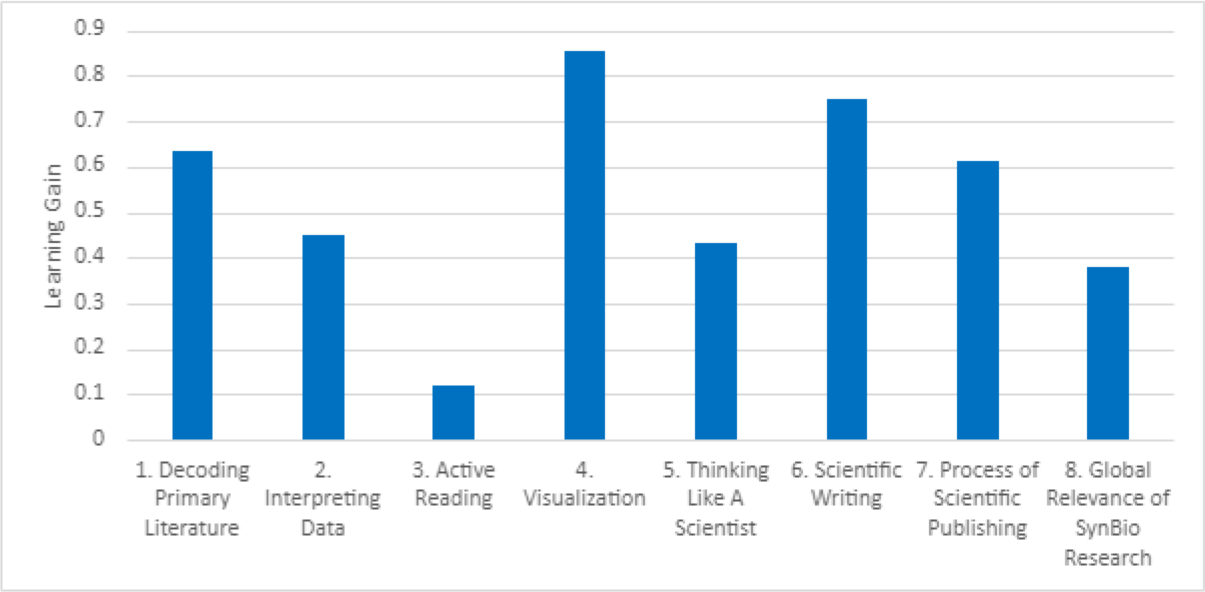

**Supplementary Figure S8. Learning gains by category**. Average post-project score minus the pre-project score for each student. n=24. Analysis includes Q7-3.

### 1.2 Supplementary Tables

#### **Supplementary Table S1.** Annotated Bibliography Student Examples.

General Instructions: All team members, regardless of the role(s) defined in the topic proposal document, will produce an annotated bibliography of at least 3 relevant articles. No two students may annotate the same article, so please coordinate your efforts! I recommend a Google Sheet to track the articles, including the text of the annotations, and who annotated them. (the link to that Google sheet can suffice as a submission).

| Student Name | Citation | Annotation |
|---|---|---|
| Student 1 | Jatain, I., Dubey, K. K., Sharma, M., Usmani, Z., Sharma, M., & Gupta, V. K. (2021). Synthetic biology potential for carbon sequestration into biocommodities. Journal of Cleaner Production, 323, 129176. https://doi.org/10.1016/j.jclepro.2021.129176 | This article reviews both natural and synthetic carbon fixation pathways that can be used to convert excess carbon dioxide in the environment to more valuable byproducts. The article also discusses strategies for the construction and optimization of novel synthetic carbon fixation pathways. While the article does not focus on plant synthetic biology, the concepts described are relatively broad and could also be applied to plant systems. Relating the methods for construction of metabolic pathways for carbon sequestration to the issue of pollution control will help us connect urban sustainability and synthetic biology in our review article. |
| Student 2 | Jia X, Bu R, Zhao T, Wu K. Sensitive and specific whole-cell biosensor for arsenic detection. Appl Environ Microbiol. 2019;85(11):1–10. https://doi.org/10.1128/AEM.00694-19 | In this article, a whole cell biosensor was designed for arsenic detection by using a positive feedback loop to address issues with sensor specificity and sensitivity. The study used E. coli to construct the biosensor as it naturally has the ars operon which can respond to arsenic. For the study, two whole cell biosensors were constructed to detect arsenic; the first biosensor did not have a positive feedback loop and the second biosensor contained a introduced positive feedback loop. The positive feedback loop uses luxR, which is regulated by the arsR-Pars regulatory circuit. When arsenic is present, luxR is activated before activating the expression of LuxR and mCherry in the Pluxl promoter. Compared to the biosensor without a positive feedback loop, the postive feedback biosensor had greater sensitivity, selectivity, and an increased output signal when arsenic is present. The arsenic detection limit of the positive feedback biosensor was also below the WHO drinking water standards. This article provides a good example of potential applications of arsenic biosensor and how biosensor design can be improved. |
| Student 3 | Harding, C.M., Nasr, M.A., Scott, N.E. et al. A platform for glycoengineering a polyvalent pneumococcal bioconjugate vaccine using E. coli as a host. Nat Commun 10, 891 (2019). https://doi.org/10.1038/s41467-019-08869-9 | This article shows the successful experimentation of a new enzyme called an O-linking oligosaccharyltransferase that can link polysaccarides to proteins capped with glucose at the end. Specifically they achieve the conjugation of the polysaccaride CPS14 to its acceptor protein ComP. The process is carried out in the bacterium E. Coli which makes this method easily accessible and reproducible for many laboratories. Meaning this technology could make cheaper processes and thereby cheaper vaccines (the current vaccines for pneumococcal diseases are expensive and labor intensive as stated in this article). The article also mentions that this enzyme can be used to link other combinations of sugars and proteins which means this technology can be used for vaccines against many diseases. |

**Supplementary Table S2. University Course Report Survey.** Text and questions of the University's standard and required student course report survey.

| You can help improve the quality of teaching at WPI by providing your responses on this form. Please consider each reply thoughtfully. These reports are used by the instructor for self-improvement, by students during course selection and by members of the administration and faculty committees. Your responses are anonymous and optional. Your comments will not be returned to your instructor until after the grading deadline. | |
|---|---|
| Question Prompt | Scale |
| 1. My overall rating of the quality of this course is | (1) Very poor<br>to<br>(5) Excellent |
| 2. My overall rating of the instructor's teaching | |
| 3. The educational value of the assigned work was | |
| 4. The instructor's organization of the course was | |
| 5. The instructor's clarity in communicating course objectives was | |
| 6. The instructor's skill in providing understandable explanations was | |
| Relative to other college courses I have taken: | |
| 7. The amount I learned from the course was | (1) Much less<br>to<br>(5) Much more |
| 8. The intellectual challenge presented by the course was | |
| 9. The instructor's personal interest in helping students learn | |
| 10. The instructor stimulated my interest in the subject matter | |
| 11. The amount of reading, homework, and other assigned work was | |
| How frequently were the following statements true in this course? | |
| 12. The instructor was well prepared to teach class. | (1) Never<br>to<br>(5) Always |
| 13. The instructor encouraged students to ask questions. | |
| 14. The instructor treated students with respect. | |
| 15. Instructor feedback on exams/assignments was timely and helpful. | |
| 16. The exams and/or evaluations were good measures of the material covered. | |
| 17. My grades were determined in a fair and impartial manner. | |
| 18. What grade do you think you will receive in this course? | (A, B, C, NR/D/F, Don't Know) |
| 19. On average, what were the total hours spent in each 7-day week OUTSIDE of formally scheduled class time in work related to this course (including studying, reading, writing, homework, rehearsal, etc.)? | (0, 1-5, 6-10, 11-15, 16-20, 21 or more) hours/week |
| Open Response Questions: Your thoughtful answers to the following questions would be helpful to your instructor. (Please answer in the space provided underneath each question.) | |
| What did you particularly LIKE about this course/lab? | |
| What did you particularly DISLIKE about this course/lab? | |
| Can you suggest anything that the instructor could do to improve the quality of teaching? | |
| Would you encourage a friend to take a course from this instructor? Why or why not? | |
